# Brooding brittle-star is a global hybrid polyploid swarm

**DOI:** 10.1101/2023.06.29.547001

**Authors:** Andrew F. Hugall, Maria Byrne, Timothy D. O’Hara

## Abstract

The widespread and abundant brooding brittle-star (*Amphipholis squamata*) is a simultaneous hermaphrodite with a complex mitochondrial phylogeography of multiple divergent overlapping mtDNA lineages and can exhibit high levels of inbreeding or clonality and unusual sperm morphology. We use exon-capture and transcriptome data to show that the nuclear genome comprises multiple (>3) divergent (π > 6%) expressed components spread across the mitochondrial lineages, and encompassing several other genera, including diploid dioecious dimorphic species. We also report a massive sperm genome size in *A. squamata*, an order of magnitude larger than in the sperm of other brittle-star species, consistent with our genetic measures of elevated and variable ploidy (>6). We propose that *A. squamata* (and related taxa) is a hybrid polyploid complex with many independent hybrid origins, variable ploidy, and complex patterns of parental subgenomes. We hypothesize that *A. squamata* has facultative sperm-dependent asexual reproduction, where sperm is required for embryogenesis but the egg and sperm only occasionally undergo fertilisation, a process that has been associated with the formation of polyploid hybrid swarms in other taxa [1]. Unique amongst known marine allopolyploids, the *A. squamata* complex inhabits an extensive bathymetric as well as geographic range. A. squamata is a much-studied animal amenable to laboratory culture: appreciating it as a hybrid polyploid complex makes it even more interesting to the study of evolutionary biology.

## 1. Introduction

Whole genome duplication (polyploidization) is a dramatic genetic rearrangement that is surprisingly well tolerated in some groups of eukaryotes [2]. It can immediately alter the fitness of the individual and, if beneficial, facilitate novel evolutionary transitions. Ancient polyploid events appear to have occurred throughout eukaryote evolution leading to diversification in the function of important gene families [2]. The formation of extant polyploids is frequently associated with hybridisation events (allopolyploidy), and their maintenance with reproductive assurance (e.g. asexuality or selfing) [3].

Research into polyploidy has focused on a few taxonomic groups, most notably angiosperms and vertebrates. In the marine environment, polyploidy has been noted in widespread asexually reproducing hermaphroditic bivalve species [4-7]. But research is almost completely lacking in phyla such as echinoderms and cnidarians which are known to exhibit varied asexual reproductive strategies [3]. The study of polyploidy in echinoderms has been hampered by practical problems in observing cytogenesis, including small chromosomes, tight clustering, low mitotic index and difficulties in obtaining meiotic preparations [8]. However, new DNA and RNA sequencing methodologies open an alternative path for research of polyploidism in echinoderms [e.g. 9].

Here we report a divergentglobal allopolyploid swarm in the common and easy-to-culture marine brittlestar *Amphipholis squamata* (Delle Chiaje, 1828 [10]) from target-enriched next-generation sequencing data. *Amphipholis squamata* (Figure 1a) is one of the most widespread and abundant benthic marine invertebrates. It has a wide tolerance of different environmental conditions. It has a nearly cosmopolitan distribution in coastal habitats, absent only from polar, brackish and abyssal environments. It has been also reported from upper bathyal habitats across Atlantic, Indian and Pacific Oceans down to below 1350 m depth [11]. This species has been called a biogeographic ‘paradox’ as it has achieved this enormous geographic range without a larval dispersal stage [12]. Instead, it releases live young (Figure 1d), generally an indicator of limited dispersal capabilities. Its success is due to its ability to raft across oceans in algal holdfasts or on other coastal debris [13, 14]. It can also coil its arms and form a ball-shape to facilitate rapid local movement to new substrata if dislodged or disturbed [15]. It is one of the first echinoderm species to colonise coastal zones around recently emerged mid-oceanic islands, such as Tristan da Cuhna [14] and Saint Paul/Amsterdam Islands [16].

**Figure 1.**
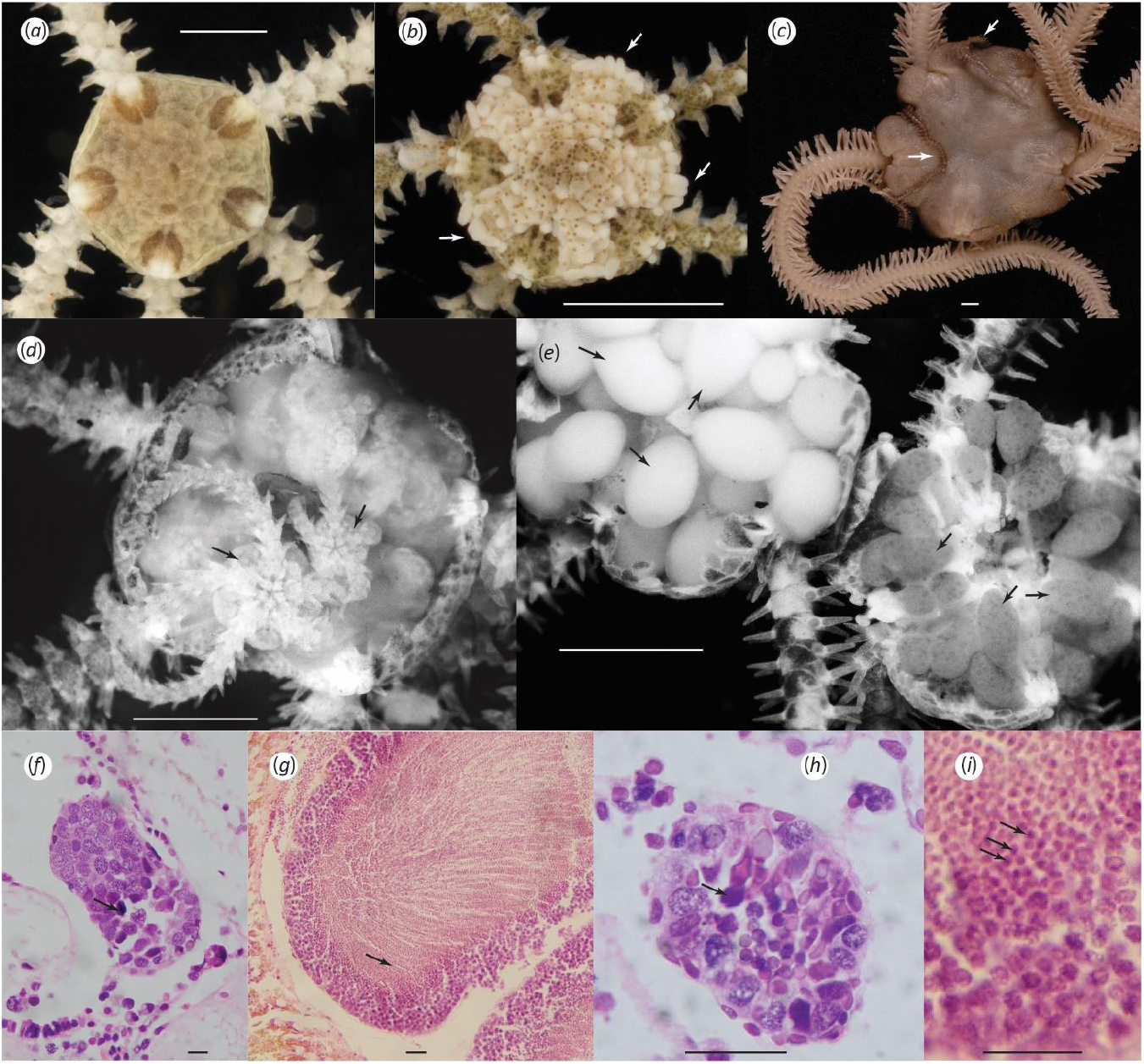
(*a*) *Amphipholis squamata*, dorsal disc and arm bases, (*b*) *Amphistigma minuta*, arrow indicates examples of the tubercle-shaped plates on the disc margin, (*c*) *Ophiodaphne formata*, arrows indicate the arms of the male emerging from underneath the disc, (*d*) *A. squamata*, dorsal disc removed to reveal brooded juveniles, (*e*) *A. pugetana*, dorsal disc removed to reveal dioecious gonads of male (upper left) and female (lower right), (*f*) *A. squamata* entire testis with only a few mature sperm (arrow) at a time, (*g*) *A. pugetana* testis with abundant sperm (arrow) dominating the lumen, (h-i) enlarged *A. squamata and A. pugetana* testis. Scale bars (*a-e*) 1 mm, (*f-i*) 0.02 mm.

The ability of *A. squamata* to successfully colonise remote locations is undoubtedly enhanced by its life history. Genetic evidence suggests a very high rate of self-fertilisation or clonal development [17] which allows colonisation of new localities from a single individual. It is a simultaneous hermaphrodite, with both male and female gametes released via gonoducts into bursal sacs at the base of each arm, where fertilisation is suggested to occur and development proceeds [18, 19]. Individuals can reproduce in isolation [19, 20]. The appearance of the sperm is highly unusual with the flagellum placed at a 70° angle to the spermatazoan axis which causes the sperm to swim eccentrically in 3-dimensional spirals while slowly rotating about its own axis [19]. Fertilisation has not been observed due to the sporadic release of single mature eggs [19]. The embryos develop from minute eggs into a vestigial pluteus larvae that remains attached to the bursal wall where they are provided with maternal nutrients via the haemal sinus [21, 22]. They metamorphose and grow into juveniles (Figure 1d) that can reach a large size as they continue to be provisioned by the parent. They eventually leave the adult through the bursal slit [18]. One egg at a time is deposited in a bursa, embryos and juveniles of various sizes can co-occur in an adult [18], but release can be seasonal in many temperate populations [23]. Adults live for 1-2.5 years in the wild [23]. Brooding can facilitate rapid local increase in abundance, and *A. squamata* can each densities exceeding 2000 animals per litre of algal matter in sheltered coastal lagoons [24].

Mitochondrial DNA forms a series of highly divergent widespread sympatric clades indicative of an ancient species-complex [12, 25-28]. COI (K2P distance) divergence has been reported to exceed 23% [29], and so most mitochondrial DNA studies have focused on the slower evolving 16S gene, where seven deep-clades (A-G) have been categorised [28, 30] (herein termed mito-groups to distinguish them from the variable phylogenies). Two of these groups were found to be congruent with limited nuclear intron/microsatellite data and considered to represent biological species [27, 28], although they were not consistent (except locally) with variable phenotypic characters based on skeletal shape, colour, or bioluminescence [26, 30, 31]. Instead, many of these mitochondrial groups are very widespread. Group A has been recorded from numerous temperate sites from the NE Atlantic, Mediterranean, both coasts of the USA, South Africa, New Zealand and Chile [30]. B has a similar range, although is encountered less frequently [30, 32], C and D have only been reported from southern Australia and New Zealand, E (and possibly G) is circum-tropical, and F only from Chile [30]..

In a program of taxonomic and biogeographic studies of the Ophiuroidea, we have sequenced ∼1500 nuclear exons (285kb) from a number of samples within, and sister to, the *A. squamata* complex. Several of these samples contained a surprisingly high rate of allelic heterogeneity [33]. Here we analyse that data and conclude that *A. squamata* is a global swarm of sympatric allopolyploid lineages, the first such complex discovered within the phylum Echinodermata.

## 2. Materials and methods

The genetic data processing used here was derived from our ophiuroid exon-capture system [33-35]. Briefly, assembled transcriptome data were used to design a set of 120 bp probes tiled to target 1496 exons in 416 genes from DNA samples sequenced using an in-solution RNA target (exon-capture) enrichment procedure and Illumina 125 and 150 bp paired-end sequencing. Raw reads were de-duplicated (Clumpify [36]), trimmed (Trimmomatic [37]) and mapped (BLAT [38]) against a special composite de novo assembled (Trinity,[39] or Tadpole [36]) sample-specific reference (see Table S1 for results and [33] for details on methods).

Our phylogenomic dataset now includes 50 transcriptomes and 1946 exon-capture samples. For this study we have included one transcriptome and 24 exon-capture samples (focused around *Amphipholis squamata* (Order Amphilepidida, Family Amphiuridae) and close relatives (Table 1, Figure 1), 12 of which were previously unpublished (supplementary table S1). Close relatives included *Amphipholis linopneusti* Stöhr, *A. pugetana* Lyman, *A. sobrina* Matsumoto, *A. torelli* Ljungman, *A*. sp2 (an undescribed species from the Kermadec Islands), *Amphistigma minuta* Clark, *Ophiodaphne formata* (Koehler), *O. scripta* (Koehler), *O. materna* Koehler, and *Ophiosphaera insignis* Brock. We selected the type species of *Amphipholis, A januarii* Ljungman, and a related species of *Amphioplus, A. depressus* (Ljungman), as outgroups based on phylogenetic similarity and good read coverage. The tissue samples were sourced from ethanol-preserved museum specimens (Table S1). Brooded juveniles and gonads were removed where observed.

**Table 1.** Description of samples and summary of sequence variant analyses. Seq vars=sequence variants, NML=Non-monophyletic lineages

| Genus | Species | Sample | Locality | mito group | Single read (110 bp) analyses |  |  |  | Merged read (190 bp) analyses |  |  |  |
| --- | --- | --- | --- | --- | --- | --- | --- | --- | --- | --- | --- | --- |
|  |  |  |  |  | No. exons | Read coverage | Median no. seq vars | Proportion of exons with > 2 seq vars | No. exons | Median nucleotide diversity | Median maximum divergence | Median NMLs |
| *Amphipholis* | *squamata* | F211339 | Australia, Victoria, Anglesea, 0-1 m | A | 360 | 221 | 4 | 0.82 | 197 | 0.048 | 0.068 | 3 |
| *Amphipholis* | *squamata* | F254622 | Australia, Victoria, Gabo Island, 0-1 m | A | 358 | 1359 | 4 | 0.82 | 200 | 0.046 | 0.068 | 3 |
| *Amphipholis* | *squamata* | MRG808.F222771 | Australia, Victoria, Port Welshpool, 0-1 m | A | 350 | 102 | 4 | 0.81 | 15 | 0.040 | 0.053 | 3 |
| *Amphistigma* | *minuta* | F173962 | Australia, Victoria, Flinders, 0-1 m | A | 304 | 70 | 3 | 0.72 | 104 | 0.052 | 0.075 | 2 |
| *Amphistigma* | *minuta* | F211345.2 | Australia, Victoria, Flinders, 0-1 m | A | 358 | 163 | 3 | 0.77 | 183 | 0.061 | 0.088 | 2 |
| *Amphipholis* | *squamata* | GLB.009 | Australia, Victoria, Lakes Entrance, 1-2 m | A | 158 | 56 | 3 | 0.75 | 22 | 0.029 | 0.039 | 3 |
| *Amphipholis* | *squamata* | A65682 | South Africa, Moullie Point, 0-2 m | A | 314 | 69 | 3 | 0.72 | 97 | 0.033 | 0.046 | 3 |
| *Amphipholis* | *squamata* | DZMB53422C | South of Iceland, 1588 m | B | 282 | 48 | 3 | 0.58 | 79 | 0.028 | 0.039 | 2 |
| *Amphipholis* | *squamata* | DZMB55448B | South of Iceland, 214 m | B | 359 | 373 | 3 | 0.65 | 197 | 0.034 | 0.045 | 3 |
| *Amphipholis* | *squamata* | IE.2013.10780 | New Caledonia, Ile des Pins, 493-694 m | C | 359 | 679 | 1 | 0.03 | 199 | 0.012 | 0.012 | 1 |
| *Amphipholis* | *squamata* | Misaki029 | Japan, Sagami Bay, 109-160 m | C | 360 | 743 | 3 | 0.5 | 200 | 0.027 | 0.037 | 2 |
| *Amphipholis* | *sobrina* | Sagami39 | Japan, Sagami Bay, 218-318 m | C | 348 | 361 | 2 | 0.42 | 164 | 0.021 | 0.023 | 1 |
| *Amphipholis* | *squamata* | Misaki011 | Japan, Misaki, 0-1 m | E | 358 | 1204 | 3 | 0.68 | 200 | 0.029 | 0.042 | 2 |
| *Amphipholis* | *squamata* | MVF214040 | USA, Florida, Blue Heron Bridge, 1-5 m | E | 89 | 148 | 3 | 0.76 | 26 | 0.025 | 0.036 | 2 |
| *Ophiodaphne* | *formata* | F111999 | Australia, Ningaloo, 165-166 m | S | 345 | 102 | 1 | 0.04 | 172 | 0 | 0 | 1 |
| *Ophiodaphne* | *materna* | IE.2007.3135 | New Caledonia, Munida Seamount, 233 m | S | 353 | 119 | 1 | 0.02 | 150 | 0 | 0 | 1 |
| *Ophiodaphne* | *scripta* | IE.2007.5337 | New Caledonia, Capel Bank, 285-545 m | S | 349 | 113 | 1 | 0.02 | 178 | 0 | 0 | 1 |
| *Ophiosphaera* | *insignis* | IE.2007.7571 | New Caledonia, Chesterfield Plateau, 345-377 m | S | 332 | 85 | 1 | 0.02 | 155 | 0 | 0 | 1 |
| *Amphipholis* | *linopneusti* | IE.2007.7480 | New Caledonia, Chesterfield Plateau, 519-522 m | S | 206 | 56 | 1 | 0.02 | 86 | 0 | 0 | 1 |
| *Amphipholis* | *sp2* | AM J24929.2 | New Zealand, Raoul Is, 21 m | S | 177 | 57 | 2 | 0.16 | 87 | 0.013 | 0.013 | 1 |
| *Amphipholis* | *pugetana* | SIO E7449 | Unites States, California, Redondo Knoll, 575 m | X | 352 | 208 | 1 | 0.02 | 191 | 0.011 | 0.011 | 1 |
| *Amphipholis* | *pugetana* | SIO E7946-2 | Unites States, California, Hancock Bank, 455 m | X | 354 | 665 | 1 | 0.02 | 195 | 0 | 0 | 1 |
| *Amphipholis* | *torelli* | DZMB49260B | Faroe Island Ridge, 400 m | X | 340 | 871 | 1 | 0.01 | 185 | 0 | 0 | 1 |
| *Amphioplus* | *depressus* | KGR23021 | Australia, W Australia, King George River, 37 m | X | 296 | 153 | 1 | 0.01 | 137 | 0 | 0 | 1 |
| *Amphipholis* | *januarii* | UF11904 | French Antilles, St. Martin, Caye Verte, 1-9 m | X | 186 | 36 | 1 | 0.01 | 41 | 0 | 0 | 1 |

Our probe target also included 1431 bp of the mitochondrial COI gene. However, as mitochondrial 16S (rather than COI) has been used previously to categorise *Amphipholis* lineages, we also assembled partial fragments of this gene from transcriptomes and off-target reads for the exon-capture samples. Where we failed to obtain a sufficient length of 16S, we preformed additional Sanger sequencing using universal primers 16Sar and 16sbr [40] to obtain approximately 560 bp from the 3′ end.

### Assessing within genome diversity

To avoid the complications of attempting to assemble highly heterozygous genomes from short reads, we assessed allelic diversity directly from the Illumina paired-end reads as sequence variants, using a custom pipeline (RHACK3, see Dryad) developed from our exon-capture system [26]. Reads were pre-processed (as above) and mapped using BLAT [38] software with relaxed settings (allowing matching of up to 10% sequence difference) against a composite sample-specific reference (Misaki011, IE.2013.10780 & DZMB49260B, Table 1).

We ran two series of analyses using 1) single reads and 2) merged reads formed from pairs where left and right reads sufficiently overlapped (using FLASH software [41]). The first dataset was used for analyses demonstrating sequence variant richness and divergence, the second to achieve greater resolution in certain phylogenetic analyses. To determine sequence variants, we defined a fixed consistent window of 110 bp (single reads) and 190 bp (merged reads) near the centre of the exons (where coverage is typically greatest). Only reads that filled >80% of this window were counted, and only exons with total coverage >40 per sample were considered (>30 for merged reads). Reads were then grouped into bins that allowed 1 base difference (for read error) with the most abundant base per site chosen to represent the bin consensus (cf UCLUST consensus procedure [42]). This process provided aligned datasets of all sequence variants of all samples per exon. Out of our 1496 exons, there were 1255 exons of sufficient length to encompass a 110 bp window and 411 exons for a 190 bp window. This was reduced to 360 and 200 exons respectively (in 194/128 genes) that occurred in at least half the samples, including at least one of each mito-group (Table 1). Average pairwise p-distance (nucleotide diversity, π) and median maximum p-distance was used to measure sequence variant diversity.

The longer merged-read datasets were used to generate UPGMA p-distance dendrograms (using PHYLIP v3.695 [43]) as estimates of sequence variant phylogeny (the limited number of sites not warranting a more complex method). Firstly, we used these trees to demonstrate the diversity of parental lineages possibly contributing to a hybrid genomic complex [44]. We generated these exon dendrograms from all the sequence variants across all samples (excluding a few exons where the ingroup and outgroup samples were not reciprocally monophyletic). We then calculated the number of non-monophyletic clades of each mito-group that were formed from sequence variants for each sample in each exon tree, i.e., the number of lineages that were separated by sequence variants from other mito-groups. For example, the 3 sequence variants from misaki029 in exon ZGC73290 (Figure 2) are all separated in the phylogeny by sequences from other mito-groups and so form three lineages that are non-monophyletic. We then calculated the median number of these lineages for each sample across all exons, which we term the number of non-monophyletic lineages (NMLs, Table 1). These UPGMA trees were also used to calculate the minimum p-distance between sequence variants belonging to different mito-groups, including variants shared across groups (p-distance = 0), as additional measures of potential hybridization between the highly distinct major mitochondrial clades.

**Figure 2.**
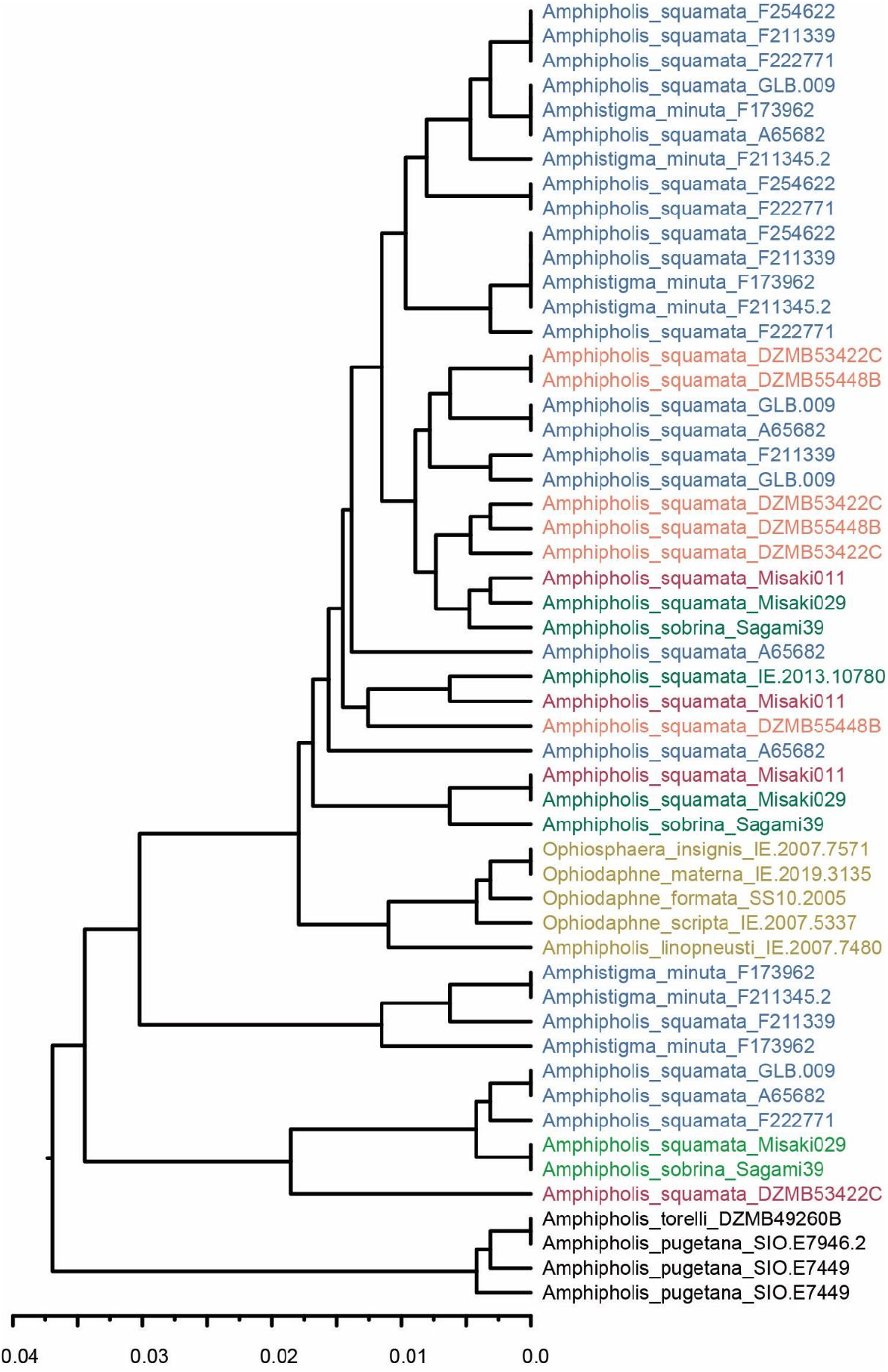
Example of an exon sequence variant phylogeny. UPGMA p-distance tree of a merged-read exon (ZGC73290 exon 1). Colour-coded by mito-group (as Figure 3). Note some samples missing due to inadequate coverage.

To investigate actual ploidy more directly, we also generated a minor state frequency (MSF) distribution (profile) [45] for each sample, calculated as the ratio of read coverage of the second most common base at a site divided by the total read coverage, for all multi-state sites across the 1496 exons with coverage >40 and less than the 95^th^ percentile of coverage across all sites (to exclude potential high copy number pseudogene artefacts). The modal shape of the MSF spectrum gives some inference on the copy number of sub-genomes, despite some data noise from vagaries of gene expression, exon-capture efficiency, and permissive base mapping [45]. A modal peak on the MSF spectrum near 50% is consistent with expectations of diploid, a peak at 33% with a triploid, and so on [45].

### Summary phylogenetic trees

We generated Maximum Likelihood phylogenetic trees for various subsets of mitochondrial and nuclear gene data, using IQTree v2.2.0 software [46] with ModelFinder selected optimal model and ultrafast bootstrap support (n=1000). Mitochondrial data comprised 1) 16S (combined GenBank accessions and our samples) aligned with MAFFT [47], and 2) combined 16S and COI for our samples only (1431 sites COI, 1314 sites 16S). Nuclear data comprised concatenated consensus (IUPAC coded) sequences, filtered by level of polymorphic sites and data incompleteness to 1325 exons in 416 genes (252.2 kb), and run with codon position optimal model. This exon data was also used to generate 234 separate gene trees (using a simple TN93 model) then summarized in ASTRAL II (v5.5.10) [17] with local posterior support values [19].

To provide another way of summarizing the genomic similarity or relatedness between samples, we used the sequence variant UPGMA trees to define a matrix of presence/absence of sequence variant lineages, defined as 1/3 of the total tree height. For this, we used the best sampled 70 exons (to minimize missing data). These presence/absence matrices were then used to generate NJ dendrograms and nMDS plots (using metaMDS() in the R library vegan, [48]).

### Flow cytometric estimation of genome size

Five ophiuroid species *Amphipholis squamata* (not sequenced, but presumably clade A or E based on its location), *Ophionereis schayeri* (Müller & Troschel), *Macrophiothrix spongicola* (Stimpson), *Clarkcoma pulchra* (Clark) and *Ophiactis resiliens* Lyman and one sea urchin *Heliocidaris tuberculata* (Lamarck) were collected from around or under boulders or algal turf from Little Bay Sydney on snorkel in shallow water (1-2 m depth). The testes of *Amphipholis* were gently detached from their attachment to the genital plates. As the testes were small (∼ 100 µm diameter) and sperm were not evident on visual inspection, they were placed on a microscope slide and checked microscopically for the presence of sperm. The sperm were gently teased out of the testes using microneedles and a micropipette was used to collect the sperm. Each testis has only ∼ 1000 sperm and so this process was repeated with 30 isolated testes dissected from 20 *Amphipholis* specimens. These samples were pooled to isolate sufficient sperm for analysis. The other species had large testes filled with sperm which oozed out. The sperm of these species were collected using a glass pipette placed in a tube and kept dry at 4°C until used for flow cytometry. The sperm of *H. tuberculata* was collected from the aboral surface of the test following injection of 0.5 M KCl, and also stored dry until use.

The absolute quantity of DNA per cell in picograms was estimated for the sperm of the five ophiuroid species by flow cytometry with an ICP 22A (Ortho Instruments). Sperm solutions were prepared by adding a few µl of dry sperm to 5 ml of 0.9% tri-sodium citrate. The fluorescence intensity in picograms of DNA per channel was calibrated using two standards of known DNA content, with *Drosophila* (*D. melanogaster* Meigen) diploid cells used as the lower genome size standard (0.36 pg DNA) [49] and the sperm of *Heliocidaris* as upper size standard (1.05 pg DNA) [50]. A drop of each standard added to 2 ml of staining solution containing 1% Triton X-100 and 1 µg of ml-1 4’,6’-diamidino-2phenylindoel 2 HCl (DAPI) (Mannheim Boehringrer) in 0.08 M phosphate buffer pH 7.3 and used within one hour (being stable for over two hours). Drops of the *Drosophila* standard, *Heliocidaris* standard, and ophiuroid sperm were added sequentially to the cytometer for each run.

### Testis histology

For histological examination of the testes, *Amphipholis squamata* and *A. pugetana* were collected near Bamfield, Vancouver Island, British Columbia, Canada. *Amphipholis squamata* were also collected from Belize. Whole discs of *A. squamata* and dissected testes of *A. pugetana* were placed in Bouin’s fixative and processed for wax histology. The blocks were sectioned (6-7 µm thick) and the sections were stained with haematoxyln and eosin. The sectioned testes were examined with an Olympus microscope, photographed with an attached camera and sperm nucleus size was measured.

## 3. Results

### Genetic data quality

Exon coverage varied substantially (Table 1) but most samples had more than sufficient sequence variant scoring [cf. 51], with some expected reduction when merging reads (proportion merged varied between 13-88%). To minimise missing data, we used various combinations of samples and loci in different analyses, an inevitable compromise with such heterogeneous data, and relied on the consistency of major patterns to reinforce our main conclusions. Due to the inherent bias in gene expression, the transcriptome sample (*Amphipholis_squamata*_MVF214040) had fewer recovered exons, and a different mix, dominated by ribosomal protein gene exons. Our samples appear to contain very low levels of contamination judging from the near-unanimous read coverage for a single COI haplotype in all samples (>1000-fold read-coverage ratio).

In the initial standard exon-capture mapping (Table S1), almost all samples named as *Amphipholis squamata, A. sobrina* and *Amphistigma minuta* showed anomalously high proportion of polymorphic sites (>1%) compared with outgroups and related dioecious (e.g. *Ophiodaphne*) species, and several samples of *A. squamata* and *A. minuta* from SE Australia exceeded 4%. One key exception was *Amphipholis squamata* IE.2013.10780 (see below). Despite our limited sampling, we confirm previous reports of very high divergence of the major lineages with COI differences exceeding 20% (and 7% within mito-groups), matched by maximum nuclear exon differences exceeding 8%.

### Mitochondrial clade identity

We assigned a mitochondrial clade identity to each of our samples through a tree-based comparison of our 16S sequence data with published data [12, 25-27, 30] (Figure S1). For the purposes of analysis, we designated six mito groups as a means of classifying samples that share maternal genomic lineages: previously identified groups A, B, C, and E, related taxa (S) and outgroups (X). Our samples clustered with previously identified groups A-E, however, our B samples are distinct enough to be possibly recognised as their own subgroup which we designate here as B2. 16S sequences in animals we identified as *Amphistigma minuta* clustered within group A. Similarity of COI places *Amphipholis sobrina* Sagami39 within the group C (Figure 3a).

**Figure 3.**
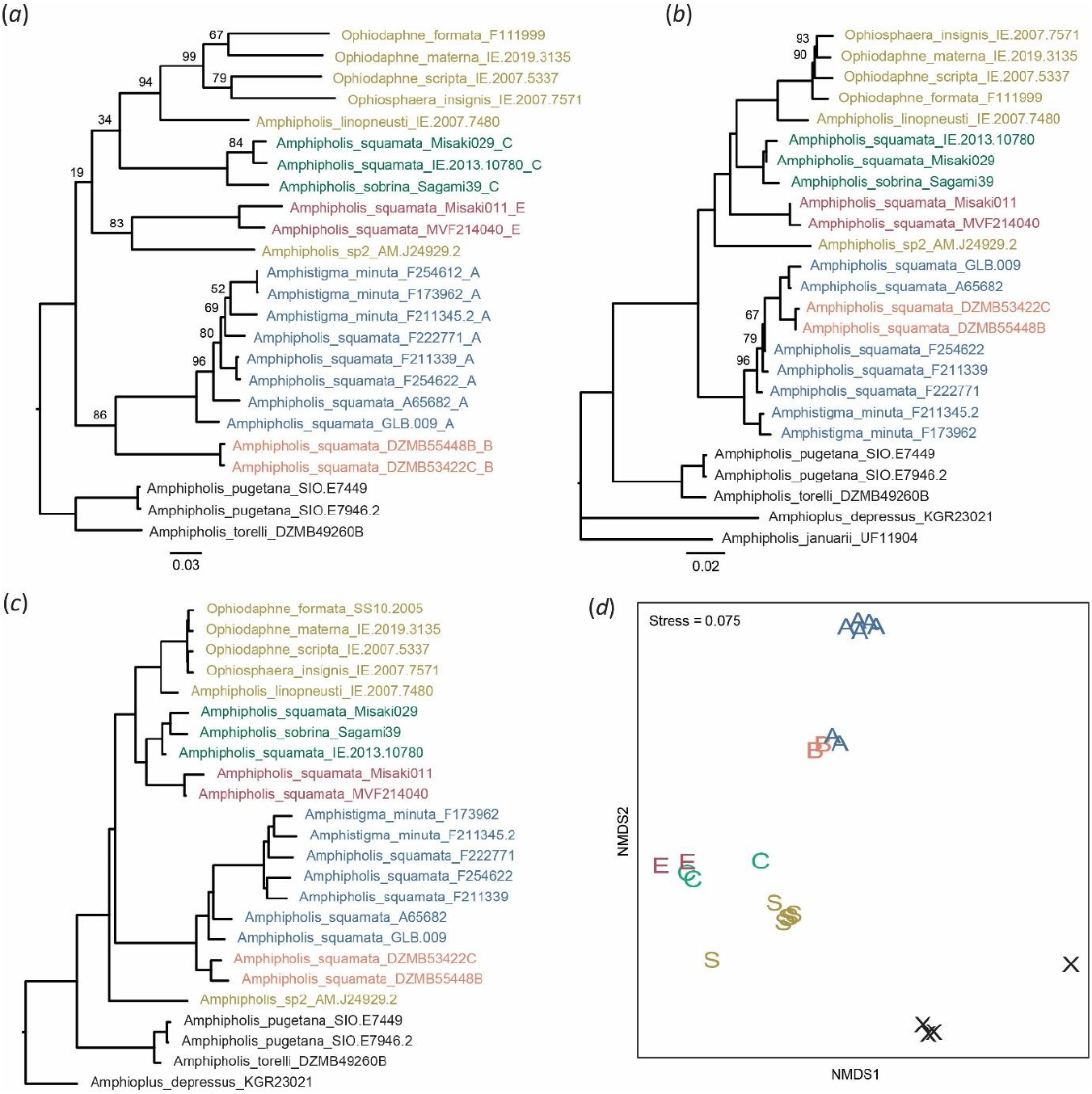
Summary phylogenetic patterns: (*a*) mitochondrial DNA (16S+COI) representing the maternal sub-genome, (*b*) standard concatenated 252 kb exon data, (*c,d*) shared sequence variant cluster lineage Jaccard pairwise distance from 70 data-rich exons, as (*c*) Neighbor-joining tree and (*d*) nMDS. Figures *b-d* provide alternative ways of representing the amalgam of parental sub-genomes. Samples are colour-coded according to *squamata* complex mito-group classification (A blue, B purple, C green, E red, S gold, black for remainder). Trees (*a*) and (b) are IQ-Tree ML with bootstrap node support. Some samples are missing from some trees due to data limitations. The ASTRAL-II tree version of (*b*) is essentially the same (Figure S2).

Our samples were generally found within known geographical and bathymetric ranges of the 16S groups. Our group A samples (including *A. minuta*) were found in southern hemisphere temperate coastal sites. Our E samples are from subtropical shallow water sites in Florida and southern Japan. Our B samples came from mid-upper bathyal sites south of Iceland which extends the bathymetric range for this group. The 1588 m sample is one of the deepest yet recorded for the whole *A. squamata* complex.

We considerably extended the known range of group C, to include outer shelf and upper bathyal (109-694 m) sites in New Caledonia and Japan. One of the C samples was identified as *A. sobrina* and our other samples of C also had this morphology (a tendency to have 4 arm spines, enlarged radial shields, distinct primary disc plates, and thinner arms compared to the typical *A. squamata*). *Amphipholis sobrina* has previously been reported from Japan and the China Sea (20-550 m) [52, 53] and goupp C predominately from 70-100 m off southern New Zealand [26] (with one specimen found in shallow water in Fiordland, a habitat known to harbour emergent deep water species [54]). In summary, group C (=*A. sobrina*) is a shelf to upper bathyal clade found in the west Pacific Ocean that occasionally occurs in shallow water. Our reported geographic range extension of groups B and C, but not A or E, reflects the sample bias of our research program, which has prioritised lesser known deep-sea environments.

### Phylogenetic analyses

While the mtDNA tree provides an estimate of the maternal lineage component of the whole genome, the combined nuclear data present a summary average of parental sub-genomes. In hybrids this can be a complex amalgam, not easily depicted as a bifurcating tree, and hence we provide several approaches to summarizing nuclear genome phylogeny.

In all phylogenies based on nuclear exon data (Figure 3b, Figure S2) we recovered two main groups, or clades, within the *A. squamata* complex: the first containing A, B and *Amphistigma minuta* and the second containing C, E, an undescribed species (sp2) from the Kermadec Islands, and a group of sexually dimorphic species (often with dwarf commensal males, Figure 1c) that are epizoic on sea urchins including *Ophiodaphne* and *Ophiosphaera* species and *Amphipholis linopneustei*, which we will refer to as the *Ophiosphaera* complex (as the earliest genus name). Excepting the A-B group, the major lineages generally have high bootstrap support. Allowing for high (saturated) divergence and limited support, the mtDNA tree is broadly similar (Figure 3a, and Figure S2) but with a key difference in the relative divergence between A and B and between C and E.

*Amphistigma minuta* was generally imbedded in the A-B clade and was only sister to this group in one of our tree analyses (concatenated exons; Figure 3b). This was a surprise as *A. minuta* specimens are easily distinguished morphologically by the presence of elevated plates around the rim of the disc (Figure 1). The A and B mito groups were only reciprocally monophyletic in the mtDNA tree (Figure 3a; Figure S1) and ASTRAL exon tree (Figure S2). Within the C-E main clade, sp2, C (=*sobrina*), E, and the *Ophiosphaera* complex formed distinct subclades, although their relationship differed between analyses.

The sexually dioecious species *Amphipholis pugetana* and hermaphroditic fissiparous species *A. torelli* formed a clade that was sister to the *A. squamata* complex. *Amphioplus depressus* and *Amphipholis januarii* were much more distant outgroups, and (along with other sequenced but not shown *Amphipholis* species such as *A. kochi* and *A. misera*) have been shown previously to be polyphyletic with respect to other amphiurid genera [34], not closely related to *A. squamata*, and a genus-level taxonomic revision is required.

### Allelic richness and diversity

The proportion of loci with more than 2 sequence variants was very high for most samples in the *A. squamata* complex (Figure 4a, Table 1; >=60%) which implies whole genome duplications or hybrid polyploidy. The median number of variants per exon per sample ranged from 1-4 (Table 1; Figure 4a, Figure S3), with the highest numbers (3-4, with up to 7 variants for some exons) occurring in the A mito group. Samples from groups B and E had slightly fewer variants (median 2-3), and C and sp2 fewer again (1-2). There was considerable variation within our group C samples, with sample Misaki029 having a median of 3, Sagami39 with 2, and IE.2013.1078 with only 1. Finally, the majority of exons in the *Ophiosphaera* complex and outgroups were homozygous, with only a few having more than 3 variants. The different sequent variant datasets were highly consistent (Figure S4).

**Figure 4.**
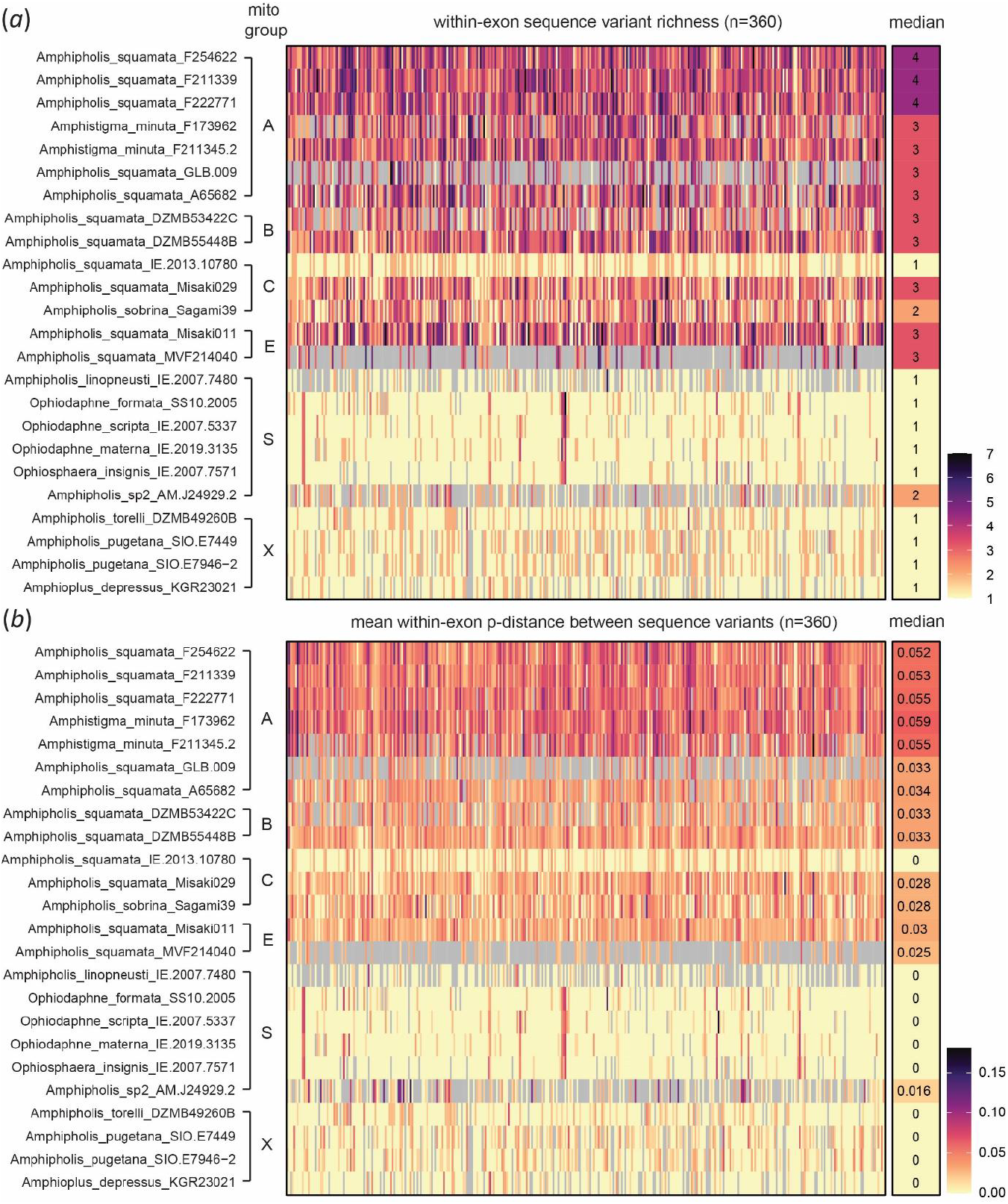
Heat-map of sample sequence variant (*a*) richness, and (*b*) mean within-exon pairwise p-distance between sequence variants from our 360-exon single-read dataset (nucleotide diversity π). Last column is the median across exons for each sample. Note the consistency between exons within a sample and between samples within mito-groups (including group E that contains a transcriptome MVF214040).

This pattern of variant richness was also reflected in variant nucleotide diversity (Table 1; Figure 4b), which ranged up to (an exceptional) 6% in the A group (including *minuta*), and 4% in C and E. Median maximum divergence across groups A-E can exceed 8%. The transcriptome (MVF214040, mito-group E) sample showed sequence variant diversity consistent with that obtained from our exon-capture E sample (Misaki011), hence these variants were expressed (transcribed as RNA), discounting bias from junk DNA pseudogenes. Again, sample IE.2013.10780 had low diversity. *Amphipholis* sp2 showed a mixed signal of elevated richness and nucleotide diversity despite some missing data. Diversity was typically low (<1%) in the *Ophiosphaera* complex and outgroups.

To investigate the origins of hybridisation, we assessed evidence for combinations of different lineages, using each sample’s mito-group as a maternal marker. Variants within a sample appear to be related to more than one mito group (e.g., Figure 2) and be scored as the number of non-monophyletic lineages (NMLs) with respect to mito-group, i.e., lineages of variants from one mito-group separated by variants from other mito-groups (Table 1, Figure S5). This provides a rough proxy for the heterogeneity of parental diversity. There was a median of 2-3 NMLs within samples in the A-B-*minuta* group and 1-2 in the C-E group. Sequence variants were generally less divergent between groups A and B, and between C and E, than for other between-group comparisons (Figure S6). Finally, the within-exon diversity also included clades of closely related sequence variants within a single mito-group (e.g., Figure 2).

Cluster and ordination of the presence-absence of sequence variants in each sample differed from the mtDNA maternal lineage phylogeny for some A and B mito group samples (Figure 3c, d). For example, samples GLB.009 and A65682 appear to be more similar to B samples than other A samples (despite the large difference in mtDNA).

### Minor Site Frequency profiles

The *Ophiosphaera* complex, our outgroup samples, and *A. squamata* sample IE.2007.10780 (mito-group C) showed peaks approaching 50% on MSF profiles (Figure 5) consistent with diploidy. In contrast, other samples from groups A, B, C & E all had peaks at <=12.5%, or a broad upward trend towards low values, consistent with at least octo-ploidy or complex aneuploid genomes. The variation in these profiles suggests potential differences in ploidy between samples. *A. squamata* IE.2007.10780 may represent a sexual parental lineage.

**Figure 5.**
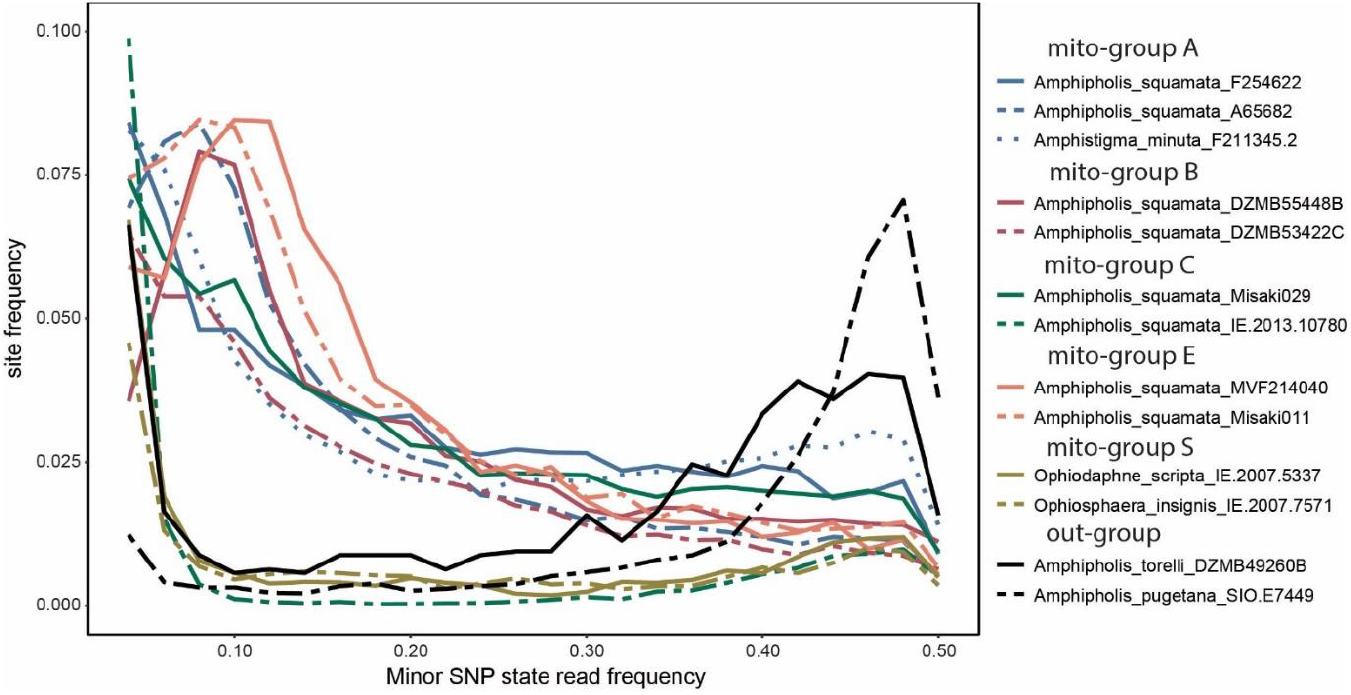
Minor state frequency (MSF) distribution of SNPs plotted in 0.02 (2%) bins for selected exon-capture and transcriptome (MVF214040) samples. Minor state is calculated as the ratio of coverage of the second most common base at a site divided by the most common for all sites with more than one base state recorded. All sites with coverage 40<>95%CI. SNP frequencies less than 0.04 not shown. MSF distributions with a peak around 0.5 (50%) conform to diploid expectations (Group S, outgroups, and the group C sample IE.2013.10780), other samples have MSF distributions indicative of elevated ploidy [45].

### Size of genome and sperm nucleus

The sperm of *Amphipholis squamata* had a genome size of 27.18 pg, approximately an order of magnitude larger than the sperm of the other species tested, including *Clarkcoma pulchra* (3.51 pg), *Ophionereis schayeri* (3.13), *Macrophiothrix spongicola* (1.80), and *Ophiactis resiliens* (2.81). These four comparison species possessed large testes and are known to have a broadcast spawning mode of reproduction [55, 56]. The sperm nucleus of *A. squamata* had a mean diameter of 5.5 µm (SE = 0.18, n = 9), while the sperm nucleus of *A. pugetana* had a mean diameter of 2.6 µm (SE = 0.08, n = 9). Measurement of the sperm nucleus in published transmission electron microscopy sections of *A. squamata* [19] also showed that they are large, 6.7 µm diameter.

## 5. Discussion

The presence of numerous sequence variants (>2) in a high proportion of our target loci (Figure 4a) is indicative of extensive genomic duplication or polyploidy occurring in the *Amphipholis squamata* complex. Both the richness of exonic sequence variants (up to 7, median 4) and minor-state frequency profiles [45] imply ploidy levels up to 8, although this maybe a conservative estimate given the limitations of our data (short next-generation raw reads). The massive genome and nucleus size recorded from the sperm of *A. squamata* compared to other species is also indicative of high ploidy. Sequence variant richness is reduced in other *A. squamata* samples (median ranging from 3 to 1) and one (IE.2013.10780) appears to be diploid. There is no evidence for polyploidy in the *Ophiosphaera* clade, *Amphipholis* sp2 and outgroups.

The presence of highly divergent nuclear sequence variants (> 8% p-distance) is indicative of hybridridisation between divergent lineages (allopolyploidy). These values are amongst the highest yet recorded [57] (once bdelliod rotifers are discounted [58]). The presence of three or more highly divergent sequence variants implies multiple hybridisation events occurring in the ancestry of these polyploids. Moreover, none of our samples are replicate clones and the presence/absence of shared alleles is highly variable between samples, also consistent with multiple hybridisation events between lineages. Ancestral duplication events with subsequent gene loss do not explain our observations of shared variants, diploid ingroup lineages and high variant divergence.

We have no direct evidence for the age of the *A. squamata* hybridisation or ploidy-elevation events. The sharing of almost identical (<= 1 bp variation) sequence variants between mito-groups A and B, and C and E, suggest relatively recent hybridisation with little subsequent divergence from mutation. However, we also have very high mitochondrial and nuclear sequence diversity (exceeding 20% K2P distance in mitochondria and >10% across our nuclear target) indicating an ancient origin for the complex as a whole. There are no reported fossils for this complex, but molecular clock estimates from published phylogenies are Paleogene in age [25][34]. This level of divergence would be considered family- or even an order-level difference in birds and mammals [59]. In addition to this deep divergence, we also found clusters of closely related nuclear sequence variants, generally restricted to within one mtDNA-clade, which could indicate post-hybridisation mutation (the Meselson effect), partial gene conversion, recombination, or inherited genetic diversity from parental lineages.

Overall, the evidence is consistent with the *A. squamata* complex being a hybrid-polyploid swarm, arising from multiple parent diploid lineages, multiple hybridisation events and multiple ploidy-elevations. Except possibly for one sample (IE.2013.10780), we lack potential diploid representatives from which we can determine the original parental lineages. Indeed, some diploid parents may no longer be extant. Consequently, we interpret our patterns of nuclear allelic inheritance in relation to patterns found in the unilineal mitochondrial DNA. Our samples are spread across four recognised mito-groups. Of these, group A (including specimens of *Amphistigma minuta*) is most closely related to B, and C (including *Amphipholis sobrina*) to E (see mtDNA trees Figure 3a and S1). The patterns of shared nuclear sequence variants reflect these relationships. Divergence between variants is generally less between A and B, and between C and E than other between-group comparisons. We interpret this as hybridisation being more likely between closer related lineages. Shared variants between A-B and C-E are much rarer and may represent low levels of homoplasy, sequence conservation or error. Sequence variant richness (here used as a surrogate for ploidy) is not consistent across the mito-groups, being highest in A, followed by B and E and then lowest but most variable in C (including the putative diploid sample).

The varying topologies and measures of similarity produced from different sequence datasets and methodologies (Figure 3, Figure S2) are indicative of varying combinations and proportions of maternal and paternal ancestral sub-genomes contained in our polyploid samples. Samples that are divergent on the mtDNA phylogeny can be brought together in our summary measures of nuclear data. For example, the group A samples A65682 and GLB.009 are closer to B group samples in the nuclear than in the mitochondrial DNA analyses, reflecting that while they share the same maternal lineage (A) these samples would have a greater proportion of B group paternal subgenomes. The distinctness of *A. minuta* and A group samples also varies between analyses, as does the relationship between C and E samples. These kinds of patterns are consistent with the combinations of sub-genomes seen in well-studied hybrid polyploid taxa [1].

Buried in the *A. squamata* complex is a lineage of putatively diploid dioecious species (the *Ophiosphaera* complex) which (for ophiuroids) exhibit unusual sexual dimorphism and host (echinoid) associations. This lineage does not seem to have participated in the *A. squamata* hybridisation events, but it is a fascinating addition to of the evolutionary complexity of the system. The diploid sister taxa we have sequenced also seem unrelated to the polyploid swarm, although, remarkably, *A. torelli* is 6-armed, asexually fissiparous, hermaphroditic, and retains its larvae within the bursae until at least the blastula stage [60]. Its sister species from the west coast of America, *A. pugetana*, is sexually diecious (Figure 1e).

Allopolyploidy is often associated with asexual reproduction such as parthenogenesis [61]. However, we do not have conclusive evidence for this in *A. squamata*. While genetic studies have repeatedly confirmed that the vast majority of *A. squamata* juveniles are identical to their mother [28] and that ‘virgin-births’ are possible [19, 20], all studies of *A. squamata* have found it to be a simultaneous hermaphrodite with paired male and female gonads. However, it is not known if the sperm of *A. squamata* have a reduced genome compared to somatic cells. The large size of the sperm of *A. squamata* compared with that of its broadcasting congener *A. pugetana*, and its large DNA content, points to the possibility of non-reduced sperm, although processes that resemble meiosis have been observed in the testes of *A. squamata* [19]. This question would need to be addressed by comparing the DNA content of the sperm and the somatic cells. The large sperm nucleus may complicate meiosis [2]. The egg cells have also been described as having relatively large nuclei [18]. In corbiculid brooding bivalves with asexual clonal-hermaphroditic reproduction the sperm are non-reduced and have the same DNA content as the somatic cells [4, 5].

Reproductive assurance [3] is also possible though selfing or sperm dependent reproduction. Observed *A. squamata* sperm have an unusual flagella that inserts into the sperm head at an oblique angle and so are not fully motile [19]. This may be a feature that promotes self-fertilisation [19]. Nevertheless, we have presented evidence for hybridisation in this paper, and other molecular studies have reported a low level of juvenile-mother divergence [62], indicating that, while clonality dominates, outcrossing must be successful in some circumstances.

Another possibility is *A. squamata* has sperm dependent asexual reproduction such as gynogenesis or androgenesis, where sperm is required for embryogenesis but the egg and sperm do not undergo fertilisation [63], as found for corbiculid bivalves. Asexuality may be facultative. Moritz & Bi [1] speculate that the formation of polyploid hybrid swarms through occasional backcrossing may be facilitated in lineages where sperm are still required for egg development.

The mechanism of allopolyploid formation varies considerably between taxa [2, 3]. We will not review all mechanisms here but note examples that share similarities with the patterns found in *A. squamata*. Polyploidy has been rarely reported from echinoderms, although this may be due to the difficulty of undertaking cytogenetic studies in this phylum [8]. Polyploid cells were reported from the echinoid *Evechinus chloroticus* but considered likely to have resulted from laboratory induced polyspermy [8]. Triploidy has been artificially induced in aquacultural holothurian species [9]. On the other hand, asexuality does occur within echinoderms via fissiparity – splitting of organisms into two pieces – in adults of ophiuroids, asteroids & holothuroids [64] and in the larvae in all classes except crinoids [65]. Many of these species also show evidence of sexual reproduction. Facultative sexual/asexual ophiuroids may represent analogues to explore the ecological and evolutionary success of facultative reproductive assurance in *A. squamata*. In the tropical species *Ophiactis savignyi* (Müller & Troschel) and *Ophiocomella ophiactoides* (Clark), sexual reproduction is rare and recruitment is mainly through cloning [66, 67]. Nevertheless, the rare sexual reproduction was seen as increasing the potential for long-range dispersal through pelagic larvae. Our genetic data suggests that *Ophiocomella* is a hybrid diploid complex [68]. Conversely, potential parthenogenisis in female-only echinoderm populations has been reported more rarely [69].

Consequently, we looked for comparative examples of allopolyploid swarms in other animal phyla. The presence of multiple hybridisations and variable ploidy occurs in well-studied *Aspidoscelis* (whiptail) lizards and *Ambystoma* salamanders, where hybrid diploids cross with sperm of other species to produce triploids and then triploids to tetraploids etc, in a mechanism that has been termed cascading polyploidy elevation [1]. This has led to a substantial diversity of both sexual and parthenogenetic lineages of varying ploidy that originated through hybridisation.

Increased ploidy, hermaphroditism, small body size, brooding and non-reductive sperm is noted for widespread bivalve species and is considered to contribute to their success [70]. *Lasaea* bivalves are a marine example of an allopolyploid swarm exhibiting variable polyploidy (3-6n), multiple hybridisation events and parthenogenisis [6, 7]. Most of its global range supports various sympatric parthenogenic/gynogenetic lineages with known planktotrophic or direct-developing diploid populations restricted geographically [71, 72]. Although analyses of mtDNA have estimated that the various *Lasaea* lineages diverged in the Neogene [72], more recent hybridisation events cannot be ruled out [7]. In the freshwater clam family Sphaeriidae, elevation of ploidy appears to be ancient, predating the Miocene-age divergence of the genera *Sphaerium* and *Musculinum*, although it is unclear whether this event included hybridisation [73].

The *Amphipholis squamata* complex shares many ecological features with these and other allopolyploids. Firstly, it is an excellent coloniser and has achieved a widespread distribution despite not having a disperse phase in its life cycle. Like *Lasaea* [6], shallow-water populations appear adapted to long distance transport through rafting, and population establishment can occur from a single individual through reproductive assurance. Consequently, polyploid lineages often have a more extensive geographic range than their diploid progenitors [74]. In *A. squamata* polyploids occur throughout the wide geographic range but it is unclear where the potential diploid lineages might be found.

Secondly, polyploid ranges frequently include more marginal habitat or environments characterised by environmental stress, where the ecological versatility conferred through hybridisation-driven allelic diversity overcomes the natural advantages of sexuality [74, 75]. *Amphipholis squamata* is most abundant in coastal habitats which experience a high degree of environmental variability and disturbance from oceanographic and climatic events. Moreover, environmental conditions vary considerably over its immense geographic range. *Amphipholis squamata* putatively exhibits enhanced fitness to climate variability through inter-population variability in life history output and seasonality [24]. Variation in colour and bioluminescence have been found to assist in predator avoidance [76]. Finally, although asexuals are in general thought to be less resistant to co-evolving parasites [77], allopolyploid swarms may offer the potential to resist localised parasitic communities. *Amphipholis squamata* is known to harbour a number of parasites, including copepods, orthonectids, turbellarians, and polychaetes [78, 79]. However, parasite presence and prevalence has been found to vary with location and colour variety [78], in line with the ecological flexibility afforded by a diverse genetic arsenal.

*Amphipholis squamata* does differ from other known marine polyploids examples in its extensive depth distribution (0-1600 m). Depth-based genetic divergence and speciation is common amongst marine lineages [80], and here we document distinct bathymetric ranges for some of the mito-groups. However, the deeper clades have been occasionally collected in relatively shallow water and the possibility exists for hybridisation to occur between deep and shallow lineages, resulting in additional ecological flexibility. Environmental and biotic characteristics that vary with depth include temperature, salinity, oxygen, light, food quality and quantity, water movement, dispersal mechanisms, host organisms, and competitor, predator and parasite communities [81, 82].

### Future directions

*Amphipholis squamata* has the potential to become a useful model organism to study the evolutionary and ecological advantages of polyploidy, hybridisation and reproductive assurance in a global marine system. It is easy to collect, culture and propagate in aquaria. One of the main goals of this paper is to promote future research into this fascinating complex.

We are currently unsure if individuals of *A. squamata* self-fertilise, are parthenogenetic, or utilise some sort of sperm-mediated strategy. This has been hindered historically by the difficultly of obtaining ripe eggs [18] and useful cytogenetic preparations [8]. Unfortunately isolating the chromosomes has not been successful for echinoderms. To understand the role that specialised asexual/clonal reproduction has played in the remarkable success of *A. squamata*, the first step would be to compare the DNA content of the sperm and somatic cells to determine if meiosis plays in reducing some or all sperm DNA content. A good understanding of its genetic architecture and life history will improve our ability to better understand unusual patterns of population genetics [28].

We also need a larger number and better geographic coverage of samples. Our current data does not fully cover the known mtDNA diversity of this complex (which is almost certainly an underestimate of the true diversity). Although, we already we have one strong candidate out of 14 *squamata-sobrina-minuta* samples, we do not know where other potential diploid ancestral populations occur, or if they even exist. Using other hybrid complexes as a guide they will be at much lower abundance or have a restricted range [83]. We do not have an adequate understanding of the geographic and bathymetric range of the polyploid lineages, nor have we yet found any obvious diploid-hybrid intermediates.

Our exon-capture system has proved useful for detecting unusual genetic signatures in species across the entire class Ophiuroidea. Potentially this could be extended to scan other little-researched echinoderm taxa, as a way of identifying targets for the more challenging life history studies. Not only would this improve our knowledge of the extent of atypical reproductive processes, but also greatly aid our understanding of the population genetics and community ecology of these species.

However, to better measure genome diversity and subgenome origins, we need longer genetic fragments. Short next-generation reads cannot be easily phased or assembled with such high levels of polyploidy. We have extracted much interesting data from our exon-capture next-generation samples, but its precision is limited for ongoing research on *A. squamata*. Long-reads from third-generation sequencers [e.g. 58] is required to better understand divergence and relationships between the numerous homologous or homeologous alleles.

## Author contributions

A.F.H., T.O’H and M.B. designed the project, T.O’H. obtained the samples and supervised the sequencing, A.F.H. performed the phylogenetic analyses, M.B. supervised the flow cytometry and histology. All authors contributed to writing the text.

## Acknowledgments

We thank Gustav Paulay, Masanori Okanishi, and Marine Research Group of Victoria for assistance in collecting specimens; the Muséum national d’Histoire naturelle Paris (MNHN), South African Museum (SAMC), University of Florida (UF), Australian Museum (AM), CSIRO, Scripps Institution of Oceanography (SIO), and Deutsches Zentrum für Marine Biodiversitätsforschung (DZMB) for providing tissue samples; Sumitha Hunjan, Kate Naughton, Claire Keely, Alexandra Weber and Maggie Haines for extracting and sequencing DNA, Valerie Morris for assistance with flow cytometry, and Paulina Selvakumaraswamy and Sam Dowland for assistance with histology.

## Data accessibility

Table S1 and Figures S1-6 are in the electronic supplementary material. Data and scripts are located in the Dryad depository https://doi.org/10.5061/dryad.qbzkh18p9

