## Supplementary figures and images for "Brooding brittle-star is a global hybrid polyploid swarm"

### Supplemental figure 1

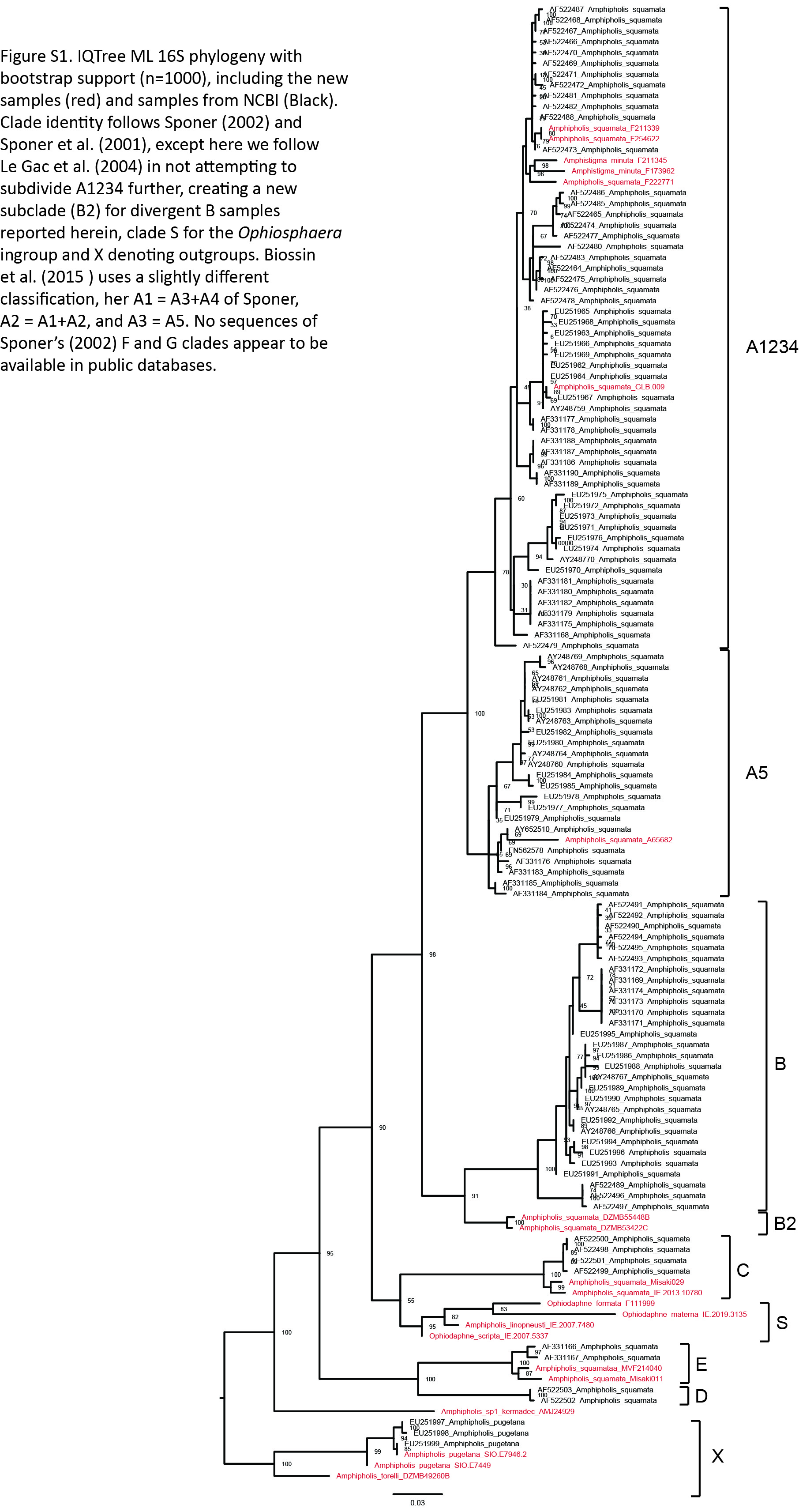

### Supplemental figure 2

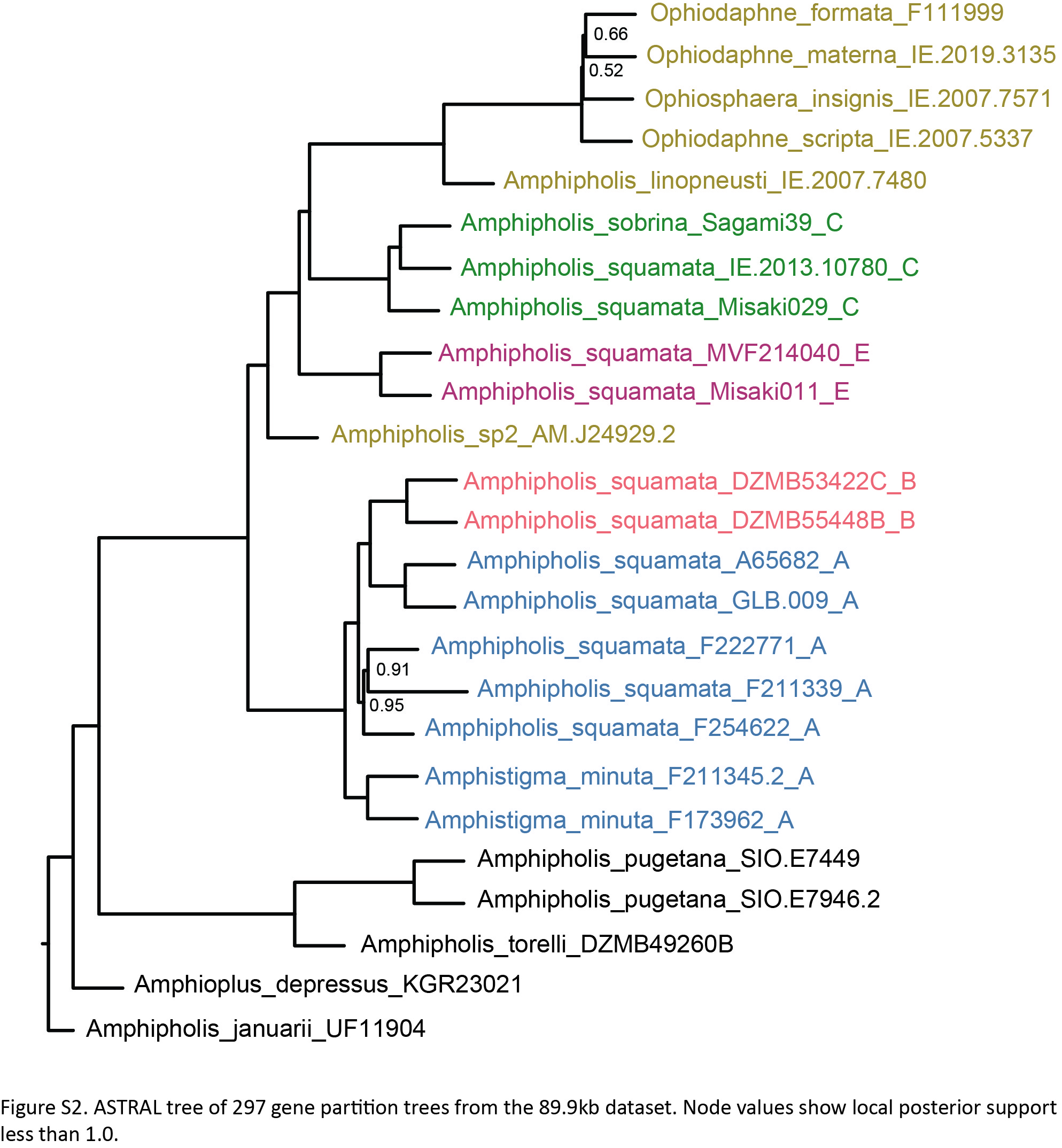

### Supplemental figure 3

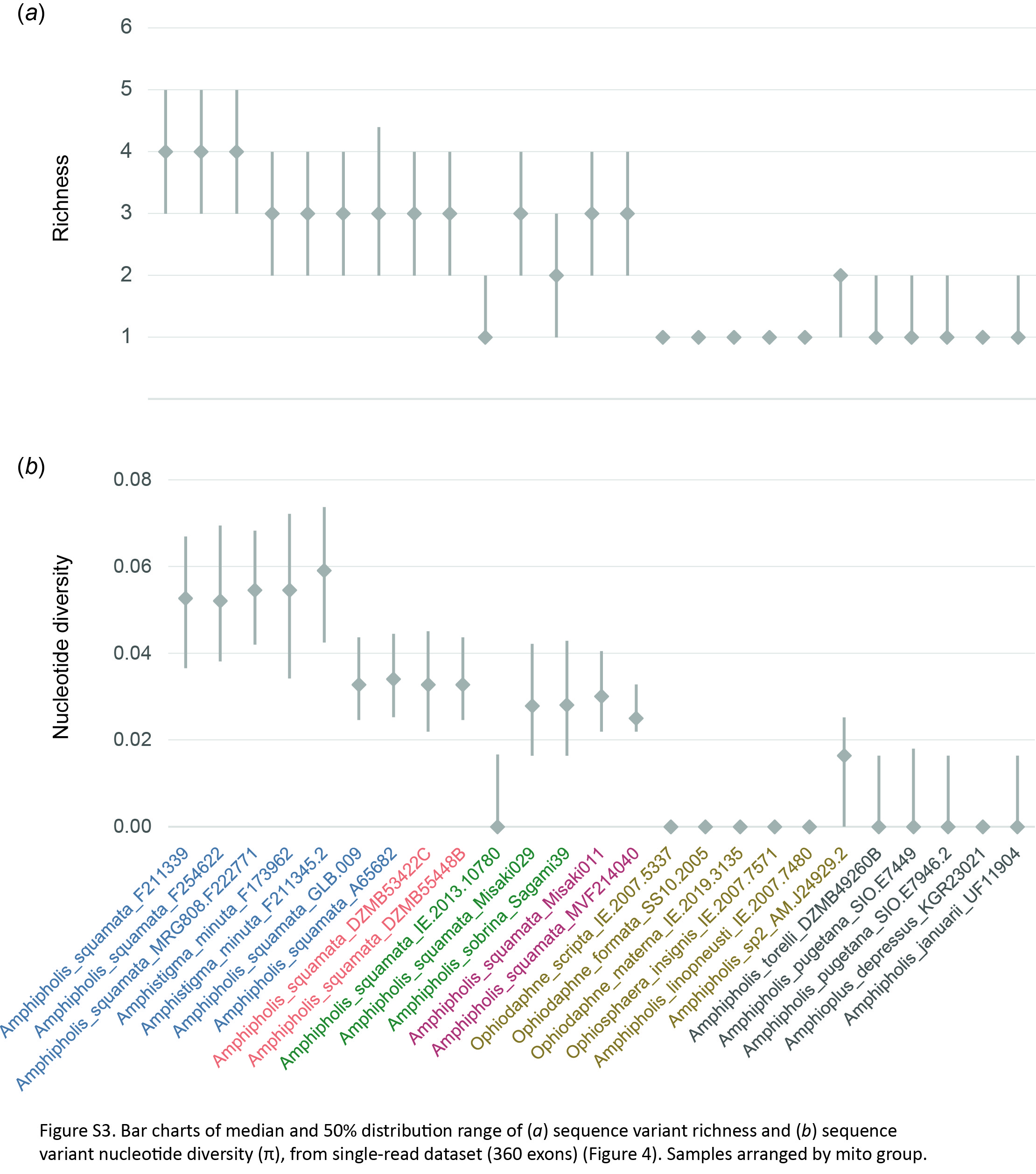

### Supplemental figure 4

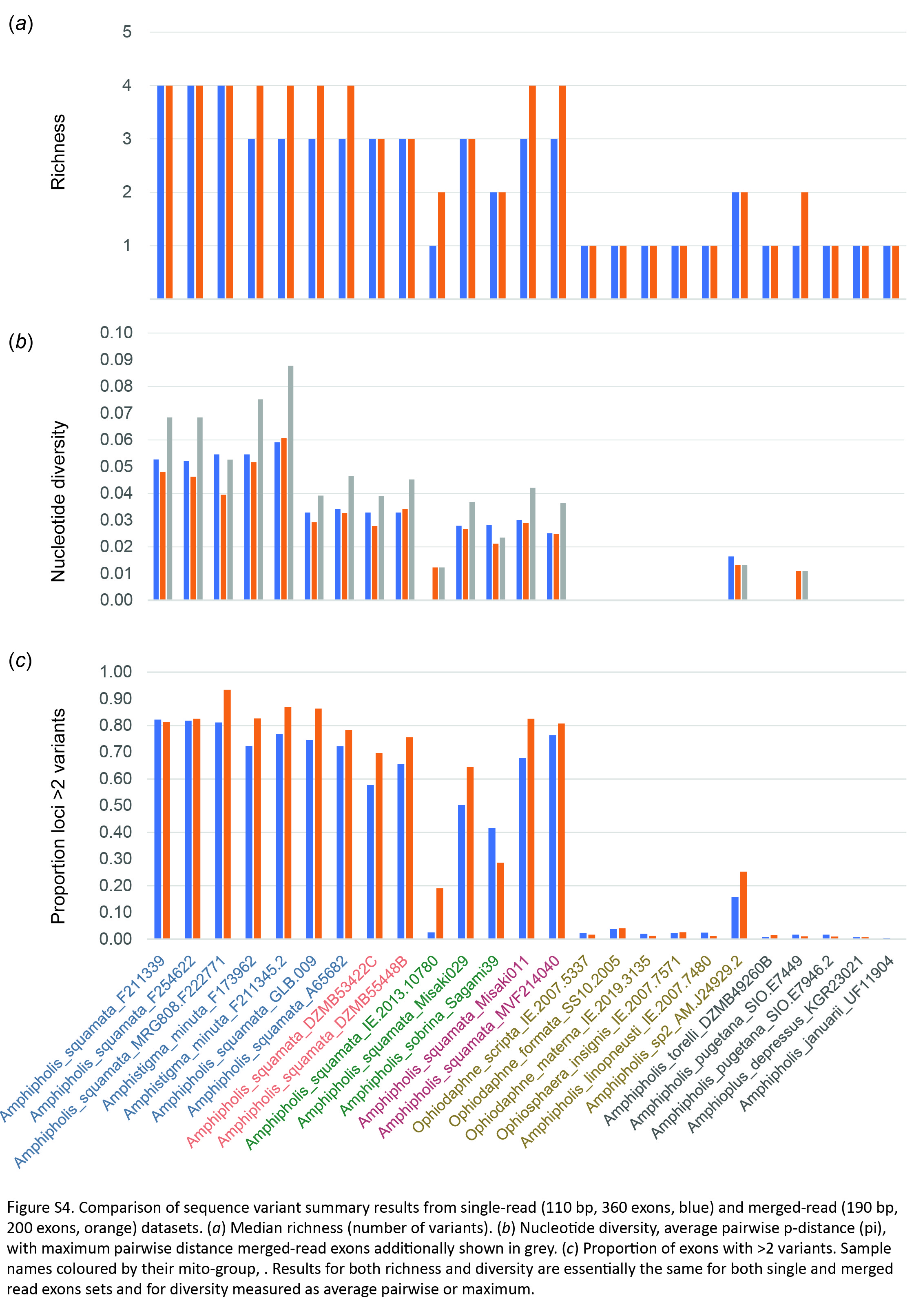

### Supplemental figure 5-6

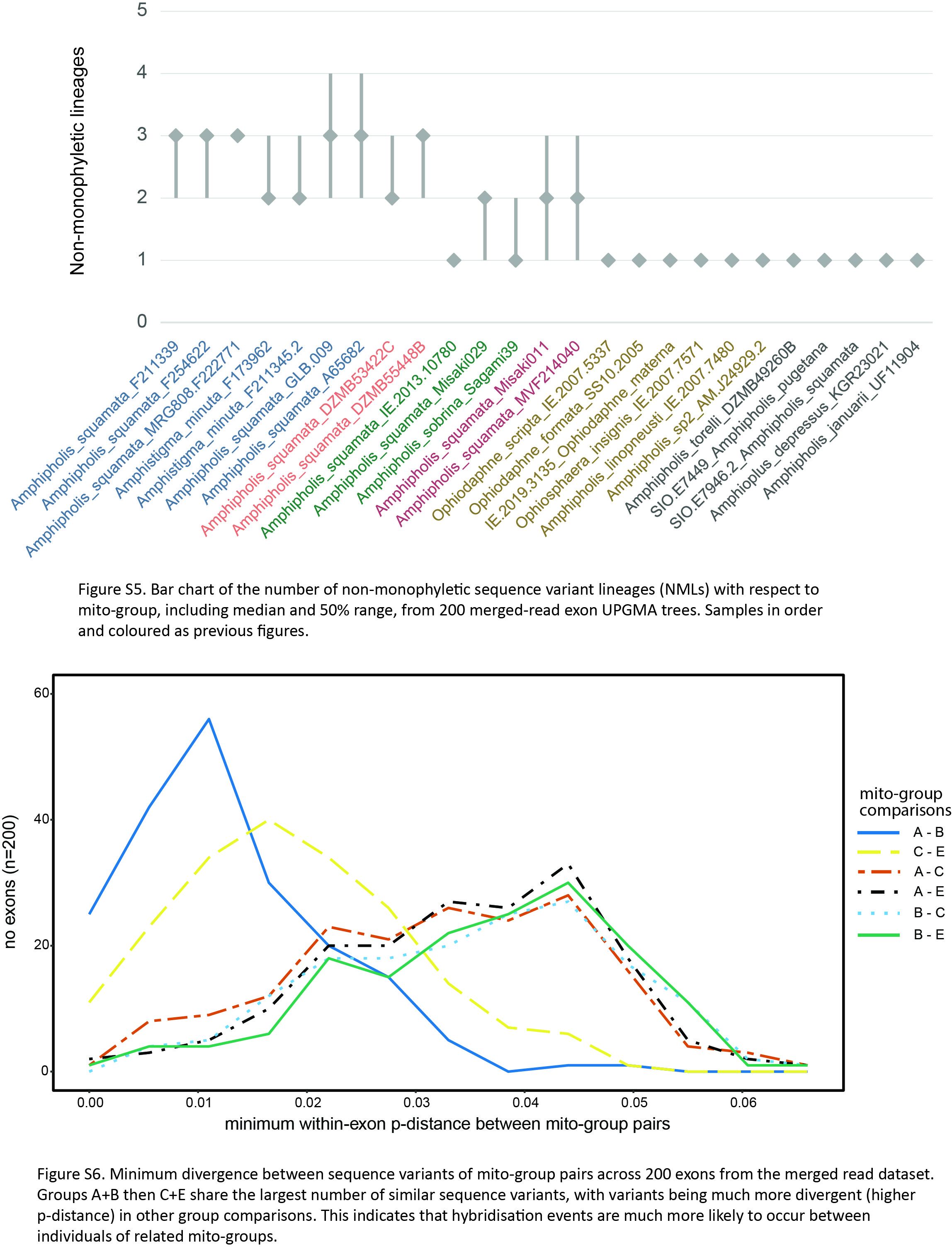
